# Effects of the anesthetic MS-222 on silver pomfret *(Pampus argenteus)* juveniles under aquaculture treatment stresses

**DOI:** 10.1101/388371

**Authors:** Na Yu, Xiaohuan Cao, Yajun Wang, Siwen Kuang, Jiabao Hu, Yang Yang, Shanliang Xu, Man Zhang, Yibo Sun, Weiwei Gu, Xiaojun Yan

## Abstract

The silver pomfret *(Pampus argenteus)* is a major economically important marine fish in China. However, *P. argenteus* is sensitive to many stress factors and susceptible to injury. This problem could be resolved using anesthesia. We determined the lowest effective dose (LED) of tricaine methanesulfonate (MS-222) and assessed the longest safe deep anesthesia time and effect after aquaculture treatment stresses. *P. argenteus* juveniles were exposed to six concentrations of MS-222 (10, 25, 50, 75, 100, and 125 mg L^-1^); LED was established at 75 mg L^-1^. The juveniles were exposed to different deep anesthesia times (4, 7, 10, 12, and 15 min) at 75 mg L^-1^; the longest safe deep anesthesia time under LED was 10 min. Finally, the juveniles were randomly divided into four groups: control group (CG), draining group (DG, drain), anesthetic group (AG, drain + MS-222 + aquaculture treatment); and non-anesthetic group (NAG, drain + aquaculture treatment). Plasma cortisol levels in the NAG, AG, DG, and CG groups were 38.739 ± 1.065 (highest), 25.083 ± 0.587, 28.644 ± 0.612, and 22.620 ± 0.836 ng mL^-1^ (lowest). The AG group showed significant differences in superoxide dismutase, catalase, and malondialdehyde activities, except for glutathione. HSP70, HSP90, GR1, and GR2 mRNA levels in the NAG group increased sharply in response to stressors. GR1 and GR2 mRNA levels in the AG group also increased significantly, whereas HSP70 and HSP90 mRNA levels showed no significant differences. Thus, MS-222 can reduce oxidative damage, stress reaction, and resistance to aquaculture treatment stresses in *P. argenteus.*

## Introduction

Anesthetic agents are widely used in modern aquaculture and have practical relevance in diverse husbandry manipulations such as grading, measurement, blood sampling, tagging, transportation, vaccination, and surgery (Wagner et al., 2015). These routine operations often affect the physiological condition and well-being of organisms and may result in undesirable reactions such as oxidative and immune stresses (Zhang et al., 2015). Under such circumstances, the use of anesthetic agents may be beneficial or even necessary. A number of chemical anesthetics have been used in fishery research, for example, MS-222, clove oil, benzocaine, and 2-phenoxyethanol (Heo and Shin, 2010) (Popovic et al., 2012; Barata et al., 2016). The most commonly used anesthetic in aquaculture practice and research is MS-222 (Popovic et al., 2012). MS-222 is a white crystalline powder that is easily dissolved in water, and rapid induction and full recovery are observed in animals. Currently, only MS-222 is licensed for use in food fish in the USA and European Union.

Cortisol is the primary glucocorticoid in teleost fish, and it is produced by the adrenal cortex or the adrenal analogue in teleost fish (the internal cells of the head kidney) (Bern, 1967). It plays an essential role in a plethora of intermediary metabolism, reproduction, and anti-inflammatory functions and stress responses (Charmandari et al., 2005). Cortisol also functions as a mineralocorticoid in teleosts, as they lack the capacity to synthesize aldosterone; thus, it is important for the maintenance of hydromineral balance (Wendelaar Bonga, 1997). The production of corticosteroids is under the control of the hypothalamus–pituitary–interrenal axis (Mommsen et al., 1999). Most glucocorticoid effects occur at the transcriptional level and are mediated by the glucocorticoid receptor (GR) (Bamberger et al., 1996; Schoneveld et al., 2004). In teleosts, two types of GRs are present: GR1 and GR2. GR1 was first identified in the rainbow trout (Ducouret et al., 1995), and cDNAs of piscine GR1 have been characterized in tilapia (Tagawa et al., 1997) and the common carp (Stolte et al., 2008). GR2 cDNAs have been characterized in the rainbow trout, common carp (Bury et al., 2003), and zebrafish (Alsop and Vijayan, 2008).

Heat shock proteins (HSPs) belong to one of the most conserved and important protein families, and they have been studied extensively as molecular chaperones with important roles in protein folding and translocation (Ming et al., 2010). HSP70, as the most conservative protein in the HSP family, plays an important role in regulating the restructure of new synthetic proteins and denatured proteins (Jiang et al., 2012). In fish, an increase in HSP70 expression not only reflects the body’s response to oxidative stress but also alleviates the adverse effects of oxidative stress with its antioxidant function (Li et al., 2014; Pierron et al., 2009). HSP90 may be involved in the cell cycle and signal transduction process and maintenance of a stable cell, and it plays an important role in the response to the stimulation of fish (Wang et al., 2004). Changes in the expression of HSP90, like HSP70, often serve as a reference index for stress in fish.

The silver pomfret *(Pampus argenteus),* a commercially important fish, has a wide distribution from the Arabian Gulf, Indian Ocean, and East Indies to Japan (Sun et al., 2017). Since the 1990s, wild-caught *P. argenteus* fisheries have suffered a severe decline because of over-fishing (Liu and Zhan, 1999) in the East China Sea. To enhance production, artificial breeding techniques have been studied in China and Kuwait (Huang et al., 2010), and studies on different aspects such as feeding behavior, breeding, growth-related genes, and health management (Shi et al., 2009a; Shi et al., 2009b; Sulaimanm and Charlesm, 2007; Azad et al., 2007) have been performed to develop culture techniques for *P. argenteus.* However, *P. argenteus* is sensitive to many types of stress factors, such as procedures for handling, grading, and transportation. Previous aquaculture studies on *P. argenteus* exposed to a transportation stressor (Zhang et al., 2017) showed that the fish panicked, leading to injury or even death. Furthermore, the fish kept swimming under normal conditions and exhibited relatively low tolerance to starvation (Liao et al., 2017). Therefore, low growth rates and high mortality have been found to be associated with the culture practices for *P. argenteus.* Reduction of stress in fish during commercial production practices is a major factor in the aquaculture industry. Currently, high concentrations of MS-222 are used for the sampling of *P. argenteus;* however, there is limited information on an appropriate concentration of MS-222 for reducing stress in this species.

Because of growing interest in the culture of *P. argenteus* and lack of detailed practical information on the administration of anesthetics, the first aim of this study was to determine the lowest effective dose (LED) on the basis of induction and recovery times of MS-222 that could be efficiently used for *P. argenteus.* The second aim was to determine the longest safe deep anesthesia time of MS-222 under LED. The effects of physiological and biochemical indicators during aquaculture treatment were also studied. Plasma cortisol levels and antioxidant capability were assessed as physiological and biochemical indicators, and the expression of stress-related genes as molecular biomarkers was analyzed to reflect the effect of each treatment.

## Material and Methods

### Fish and experimental conditions

Specimens of *P. argenteus* were collected from Xiangshan Island, Xiangshan, Ningbo in Zhejiang Province, China. No specific permission was required for the collection of *P. argenteus* from the sample sites, which were not protected areas of land. In addition, *P. argenteus* is not an endangered or protected species. Four-month-old fingerlings (250 fish; average weight, 11.2 ± 1.5 g; length, 7.5 ± 1.6 cm) were acclimated in six circular plastic tanks with a black background (capacity, 1200 L) and a constant flow of aerated seawater for two weeks before the experiment. For acclimation, the fish were fed twice a day (07:00 and 17:00) with a commercial diet (Shimonoseki, Japan) equivalent to 1.5% of their body weight, and feed was deprived 24 h before the experiment. The daily seawater exchange rate was 60%. The tanks were cleaned daily, and seawater quality parameters during the acclimatization period and throughout the experiments were as follows: seawater temperature, 29 ± 0.5 °C; pH, 7.9 ± 0.2; salinity, 30 ± 0.3 ppt; dissolved oxygen, 6.1 ± 0.6 mg L^-1^; and natural photoperiod.

### Anesthetic agent

One anesthetic agent was used: MS-222 (Canton, Shanghai). All stock solutions were prepared a few minutes before the start of each experiment; MS-222 was dissolved in seawater and added to anesthetic test tanks.

### LED of MS-222

Changes in the physiological status of the anesthetized fish were assessed in three consecutive stages for induction (I) and three stages for recovery (R), as described by Summerfelt and Smith with slight modifications based on the species-specific behavioral response of *P. argenteus* (Table 1). After two weeks of acclimation, the fish were gently scoop-netted and placed in 18 anesthetic test tanks (capacity, 20 L) containing different doses of the anesthetic solutions. The following doses of MS-222 were used: 15, 25, 50, 75, 100, and 125 mg L^-1^. Only one anesthetic dose was tested at a time. The induction and recovery times for all anesthetics were measured under the same experimental conditions by using a digital stopwatch. The seawater in each tank was aerated to facilitate complete mixing. Nine fish were used for each concentration of MS-222 (3 fish per tank; total, 54 fish). An effective dose of an anesthetic is the dose that produces general anesthesia (I3) within 3 min and allows recovery (R3) within 5 min (Mohammadi and Khara, 2015). When the *P. argenteus* specimens reached I3, they were immediately netted from the test tanks and placed on a wet surface for a period of 90 s. The duration of this period was sufficient to measure the length and weight of the fish. Subsequently, the fish were transferred to a recovery tank (capacity, 500 L) filled with fresh, aerated seawater for recording the stages of anesthesia recovery. Accordingly, LED of MS-222 was determined. After the anesthetic treatment, the fish exposed to the same dose of anesthetic were maintained in the same recovery tank to assess recovery time and recovery rate and observe behavioral changes during and after the treatment period of 7 days. The commercial feed was supplied the next morning, and the tested fish were not used again for the subsequent tests.

**Table 1.** *Stages of anaesthetic induction and recovery in Pampus argenteus*

| Stage | Behavioural response of *Pampus argenteus* |
| --- | --- |
| I1 | Loss of balance, slow swimming, partial inhibition of reactions to external stimuli |
| I2 | Erratic swimming. Fish still react to strong stimuli |
| I3 | Fish lay on the tank bottom. Slow decrease in opercular rate. No reaction to stimuli |
| I4 | Fish stop movement (over dose and or longer immersion in anaesthetic solution). Subsequent death |
| R1 | Fish still lay on bottom of the tank. Start of movement. |
| R2 | Unable to balance the head, often keeping the head up and trying to get up with tail force. Mouth tapping on the surface of the water |
| R3 | Normal swimming. Reaction to slight stimuli. |

### Longest safe deep anesthesia time under LED

After two weeks of acclimation, 45 fingerlings were randomly divided into five groups (3 tanks per group; 3 fish per tank) and placed in anesthetic test tanks (capacity, 20 L) containing 75 mg L^-1^ of MS-222. Only one time of deep anesthesia was tested at a time. The induction and recovery times for all the groups were measured under the same experimental conditions by using a digital stopwatch. The recovery time and recovery rate of the fish were recorded. When the fish reached I3, the time of deep anesthesia (5, 7, 10, 12, and 15 min) was recorded, and the fish were transferred to a recovery tank (capacity, 500 L) filled with fresh, aerated seawater for recording the stages of anesthesia recovery. After the anesthesia treatment, the fish exposed to the same time of deep anesthesia were maintained in the same recovery tank to assess the recovery time and recovery rate and observe behavioral changes during and after the treatment period of 7 days. The commercial feed was supplied the next morning, and the tested fish were not used again for the subsequent tests.

### Aquaculture treatment stresses

After two weeks of acclimation, healthy fish were randomly divided into four groups (n = 30) and placed in anesthetic test tanks (capacity, 500 L) to evaluate how different handling procedures during aquaculture treatment would affect stress responses and whether MS-222 has a stress-relieving effect on *P. argenteus.* The four treatment groups were as follows: control group (CG), draining group (DG; seawater drained off, 0.03 m^3^ min^-1^ for 10 min), anesthetic group (AG; seawater drained off like in the DG group; a high dose of MS-222 was added until the dose in the seawater was 75 mg L^-1^, the fish were chased with a PVE pipe [25 circles per min for 5 min], and seawater was added [0.06 m^3^ min^-1^, 5 min]), and non-anesthetic group (NAG, similar to the AG group, without anesthesia). Nine samples (3 tanks per group; 3 fish per tank) were obtained from each group. Blood was collected using heparinized syringes and then centrifuged at 1,500 × *g* for 15 min at 4 °C. The supernatant was transferred to a 1.5-mL centrifuge tube and stored at −80 °C until analysis. The kidney and liver were quickly removed (placed on ice) and then stored at −80 °C until subsequent analyses.

### Blood and liver parameters

#### Plasma cortisol level

Fish cortisol levels were determined using the Fish Cortisol Enzyme-linked Immunosorbent Assay Kit^®^ (Xin Yu Biotech, Shanghai), according to the manufacturer’s instructions. This kit uses the quantitative sandwich enzyme immunoassay technique. The microplate provided in this kit is pre-coated with a monoclonal antibody specific for cortisol.

#### Liver superoxide dismutase, catalase, and glutathione activities and malondialdehyde content

The liver samples were homogenized in physiological saline (1:9 dilution) and then centrifuged at 600 × *g* and 4 □ for 10 min. Superoxide dismutase (SOD), catalase (CAT), and glutathione (GSH) activities and malondialdehyde (MDA) content of the supernatant were measured using the total SOD assay kit (hydroxylamine method), CAT assay kit (visible light), reduced GSH assay kit (spectrophotometric method), and MDA assay kit (thiobarbituric acid method). These kits were purchased from Nanjing Jiancheng Biological Engineering Research Institute, China. The hepatic protein content was measured using the Micro BCA Protein Assay Kit (Beijing ComWin Biotech Co., Ltd., China).

#### Real-time quantitative polymerase chain reaction

Total RNA was extracted from the kidney tissue by using TRIzol^®^ Reagent (Invitrogen, USA), according to the manufacturer’s instructions. The RNA quality was assessed using 1% formaldehyde denaturing agarose gel electrophoresis. The purified RNA generally had an 0D260/0D280 of 1.8–2. The cDNA was synthesized using the RT-PCR Kit (TaKaRa, Japan), according to the instructions of SYBR^®^ PrimeScript™. To adjust the quantity of input cDNA, the housekeeping gene β-actin was used as the internal control. The target and reference genes used in this study were based on published information (Table 2). The stress-related genes selected were as follows: heat shock proteins (HSP90 and HSP70) and glucocorticoid receptors (GR1 and GR2). All primers were synthesized by Shanghai Biocolor BioScience & Technology Company (China). Real-time quantitative polymerase chain reaction (RT-qPCR) was performed using the Mastercycler EP Gradient Realplex (Eppendorf, Germany). SYBR Green (Roche, USA) was used as the fluorescent dye, and the manufacturer’s protocol was used. RT-PCR was performed using a total volume of 20 μL. The cycling conditions were as follows: denaturation at 95 °C for 2 min, followed by 40 cycles of denaturation at 95 °C for 15 s, annealing at 60 °C for 15 s, and extension at 72 °C for 20 s. Each qPCR was performed in triplicate, and the data for each sample were expressed relative to the expression levels of β-actin by using the 2^-▵▵CT^ method.

**Table 2.**
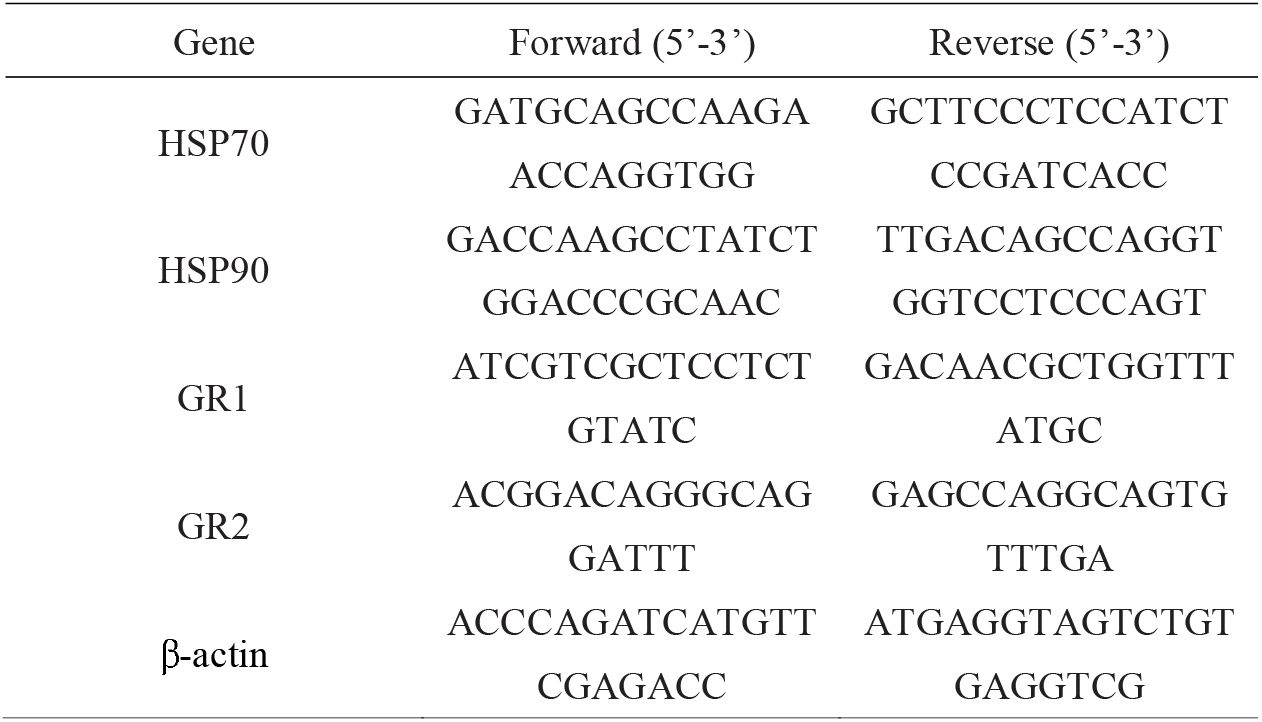
*The primers used in this study for real-time qPCR*

#### Statistical analysis

The results were presented as arithmetic mean ± standard deviation of the mean (SD) values. The data were statistically analyzed using SPSS 16.0 for Windows (SPSS Inc, Chicago, IL, USA). The relevant variables for each experimental group were compared using one-way analysis of variance, followed by the Tukey tests to detect significant differences. The minimum significance level was set at P < 0.05 in all cases.

## 3. Results

### Anesthetic efficacy

#### LED

Significant differences (P < 0.05) in the induction and recovery stages at different doses of MS-222 were detected for *P. argenteus* (Table 3). The induction times decreased significantly with increasing doses of MS-222. However, the recovery times increased with increasing doses of MS-222 (P < 0.05). At 10 and 25 mg L^-1^ of MS-222, the fish did not complete anesthetic induction and stayed at a light sedation stage (I1, Table 1). Considering the accepted efficacy criteria of general anesthetic induction time (I3) within 3 min and recovery time (R3) within 5 min, the dose of MS-222 (50 mg L^-1^) failed to induce I3 within the 3-min exposure period. The recovery time at higher concentrations (75 and 100 mg L^-1^) seemed to be similar, whereas the induction time at those concentrations was different. Furthermore, the highest concentration of 125 mg L^-1^ resulted in 50% mortality (Table 3); 75–100 mg L^-1^ of MS-222 led to no mortality during and after the treatment period of 7 days.

**Table 3.** *Induction and recovery times for Pampus argenteus anaesthetized with six doses of MS-222*

| Concentration<br>(mg L<sup>-1</sup>) | Induction (s) |  | Recovery (s) | Recovery rate (%) |
| --- | --- | --- | --- | --- |
|  | I1 | I3 | R4 |  |
| 10 | 2147.3±20.3 | - | - | 100 |
| 25 | 2400.3±15.9 | - | - | 100 |
| 50 | 1092.3±18.5<sup>a</sup> | 2275.3±103.2<sup>a</sup> | 12.3 ± 0.8<sup>a</sup> | 100 |
| 75 | 41.33 ± 4<sup>b</sup> | 167.5±50.7<sup>b</sup> | 35.5 ± 3.8<sup>b</sup> | 100 |
| 100 | 17.8 ± 3.4<sup>c</sup> | 51.7 ± 11.9<sup>c</sup> | 41.2±6.0<sup>b</sup> | 100 |
| 125 | 7.0 ± 0.9<sup>c</sup> | 19.2±2.6<sup>c</sup> | 53.3 ±4.9<sup>c</sup> | 50 |

Data are presented as mean ± SD (n=9). Statistical relationships between groups are indicated by letters where significant differences were detected ( $P < 0.05$ )

#### Longest safe deep anesthesia time under the influence of LED

The high anesthetic doses significantly reduced the time for *P. argenteus* to reach deep anesthesia. However, excessive or persistent contact with the drug can eventually lead to death. Longer the fish were exposed to MS-222, longer the recovery time needed (Table 4). After 12 and 15 min of deep anesthesia, the LED of MS-222 resulted in 78% and 33% recovery rate, respectively; therefore, 10 min was the longest safe deep anesthesia time. At the end of each exposure (4, 7, or 10 min), the fish reared in the recovery tanks quickly showed normal behavior. No mortality was observed during and after the treatment period of 7 days.

**Table 4.** Effect of LED on *Pampus argenteus* anesthetized with different deep anesthetic times

| MS-222<br>concentration<br>(mg L<sup>-1</sup>) | After fully anesthetized<br>continue anesthesia time (min) | The average time required<br>for full recovery (s) | Recovery rate<br>(%) |
| --- | --- | --- | --- |
| 75 | 4 | 43.3±6.7<sup>a</sup> | 100 |
|  | 7 | 62.5±4.5<sup>b</sup> | 100 |
|  | 10 | 182.0±4.9<sup>c</sup> | 100 |
|  | 12 | 212.7±11.8<sup>d</sup> | 78 |
|  | 15 | 249.2±15.1<sup>e</sup> | 33 |

Data are presented as mean ± SD (n=9). Statistical relationships between groups are indicated by letters where significant differences were detected (P < 0.05).

#### Plasma cortisol

Significant differences (P < 0.05) in plasma cortisol levels were observed among the groups (Fig. 1). The plasma cortisol levels of the DG, AG, and NAG groups significantly increased (P < 0.05) by 1.3-fold, 1.1-fold, and 1.7-fold, respectively, when compared with the CG group (22.620 ± 0.836 ng mL^-1^). The plasma cortisol level of the NAG group was significantly different (P < 0.05) from that of the AG group, which exhibited the highest concentration (38.739 ± 1.065 ng mL^-1^).

**Figure 1.**
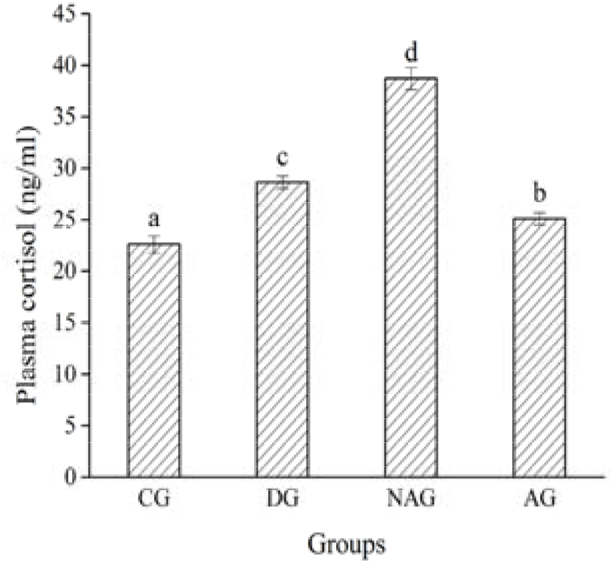
Effect of aquaculture treatment stresses on cortisol levels in plasma of juvenile silver pomfret. Data represent mean ± SD (n = 9). Statistical relationships between groups are indicated by letters where significant differences were detected (P < 0.05).

#### Oxidative stress status in the liver

The liver antioxidant capability of *P. argenteus* has been presented in Figure 2. When SOD, GSH, MDA, and CAT activities of the NAG group were compared with those of the other three groups, a significant increase (P < 0.05) was observed. However, the DG and AG groups showed no significant increase (P > 0.05) in SOD, MDA, GSH, and CAT activities. In addition, GSH values of the DG and AG groups were similar to the control values, and the CG and DG groups showed no significant increase (P > 0.05) in CAT activity. The AG group showed significantly (P < 0.05) higher GSH, SOD, CAT and MDA activities than the control before stress.

**Figure 2.**
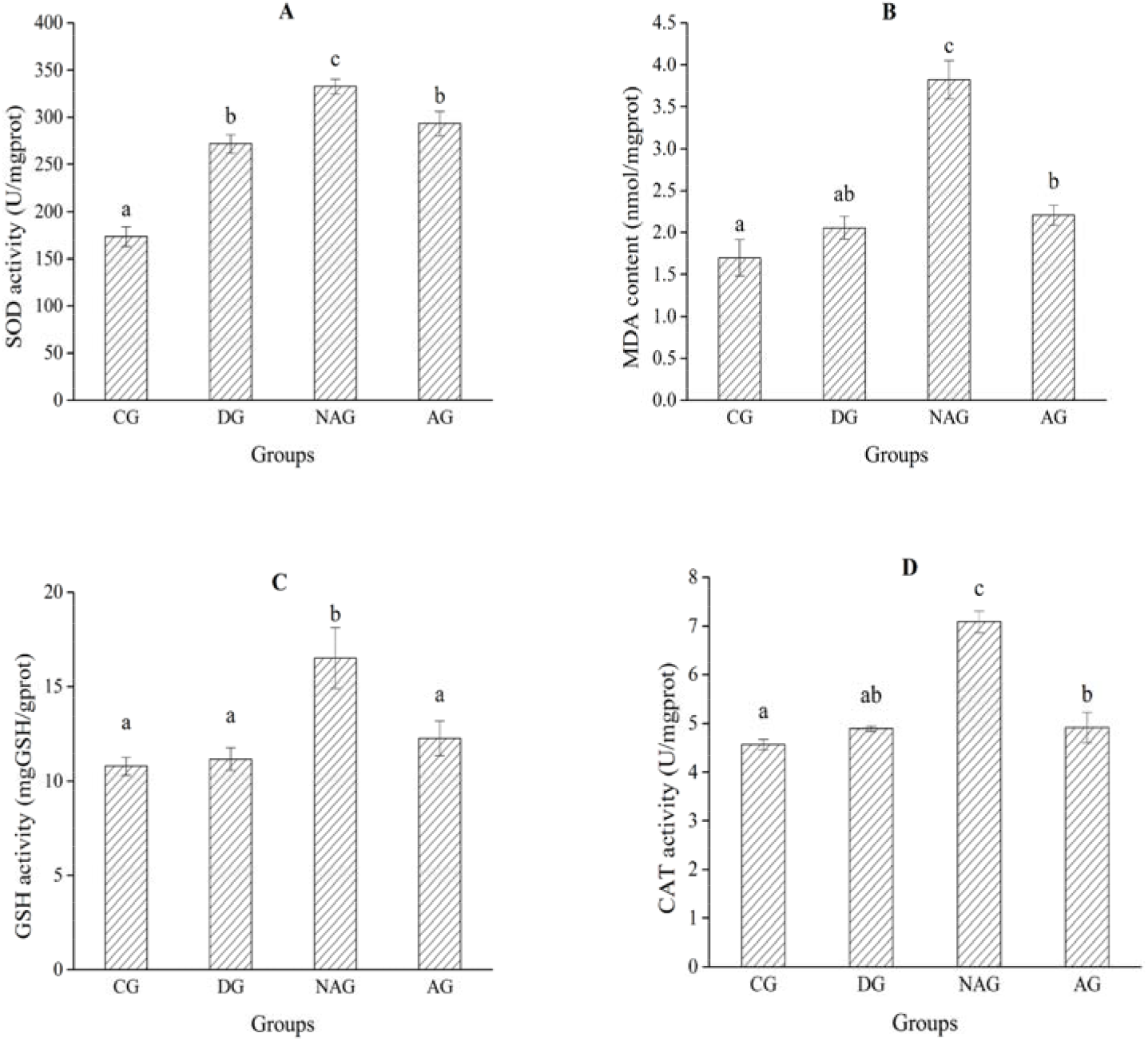
Effect of aquaculture treatment stresses on oxidative status [SOD (A), MDA (B), GSH (C), CAT (D)] in liver of juvenile silver pomfret. Data represent mean ± SD (n = 9). Statistical relationships between groups are indicated by letters where significant differences were detected (P < 0.05).

#### Stress-related gene expression in the kidney

The mRNA expression levels of HSPs (HSP70 and HSP90) and GRs (GR1 and GR2) were determined for *P. argenteus* (Fig. 3). Before aquaculture treatment, HSP70, HSP90, GR1, and GR2 mRNA levels showed no significant differences (P > 0.05), whereas HSP70, HSP90, GR1, and GR2 mRNA levels in the NAG group increased sharply. HSP70 and HSP90 mRNA levels in the AG group showed no significant differences from the CG group (P > 0.05), whereas the opposite was true for GR1 and GR2 mRNA levels.

**Figure 3.**
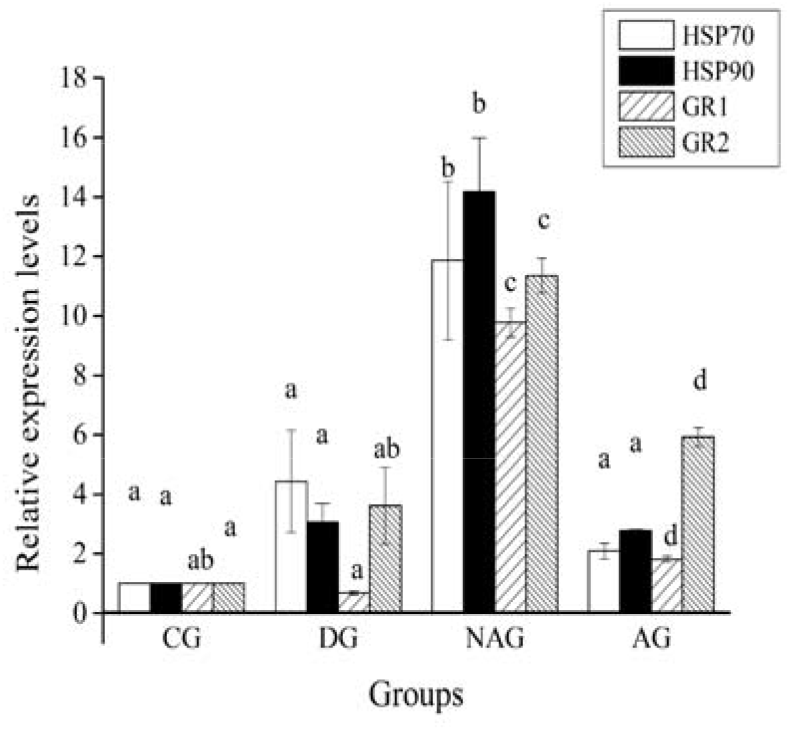
Expression levels of HSP70, HSP90, and GR1, GR2 mRNA in kidney of *P. argenteus.* The expressions of target gene were normalized by β-actin expressions. Data represent mean ± SD (n = 9). Statistical relationships between groups are indicated by letters where significant differences were detected (P < 0.05).

## Discussion

Although anesthetic agents are routinely used in aquaculture to reduce stress and physical injuries during husbandry practices, their administration and efficacy in *P. argenteus* husbandry have received scant attention. *P. argenteus* shows a strong response to stress, which can lead to death or accumulation of mechanical damage; therefore, it is important to use methods that relieve the stress of *P. argenteus.* MS-222 is particularly effective in fish because it is highly water- and lipid-soluble and readily crosses the gill membrane (Vera et al., 2010). Although MS-222 is expensive, its beneficial effects ensure its continuous application in research, food production, and ornamental fish rearing (Popovic et al., 2012). Because species may differ widely in their response to an anesthetic, it is necessary to establish appropriate concentrations of different anesthetic agents for each cultured species (Ross and Ross, 1999; Chambel et al., 2015). Factors such as body composition, sexual maturity, and gill surface to body weight ratio, which may vary with species, size, and age, may be responsible for the variations in the response (Zahl et al., 2011).

### LED and longest safe deep anesthetic time

The results demonstrate that MS-222 is an effective and safe anesthetic for *P. argenteus*. Moreover, our study defines the LED that produces desirable anesthetic effects in *P. argenteus,* namely, 75 mg L^-1^. An induction time of 3 min or less with complete recovery in 5 min is considered acceptable for fish handling (Mohammadi and Khara, 2015). This dose produced induction and recovery times of less than 3 and 5 min, respectively. In general, the induction times decreased significantly as the doses increased (Table 3). In contrast, the recovery times increased with increasing doses of MS-222 (Table 3). These results are consistent with those reported in zebrafish for MS-222 (Grush et al., 2004), whereas the recovery time for MS-222 was dose-independent in Senegalese sole (Weber et al., 2009). The explanation for these variations can be related to the fact that increasing doses are also associated with less time of contact of the fish with the anesthetic in the water, with a probable lower uptake of MS-222. Thus, when the fish is placed in recovery, the anesthetic is rapidly cleared from the bloodstream, which allows for a faster recovery (Weber et al., 2009). However, the molecular characteristics of the anesthetic, as well as the physiological and metabolic characteristics of different species, may influence the recovery time after anesthesia (Weber et al., 2011). With the prolongation of the longest safe deep anesthesia time, the fish becomes more prone to death. In the present study, we obtained a preliminary longest safe deep anesthesia time under LED. We can perform manual operations during this period to reduce the stress on the fish body.

### Plasma cortisol

Aquaculture inherently involves stressing the fish. Handling, transportation, and crowding are common stressors in fish culture, which may affect their growth, feed intake, nutrient utilization, and physiology. Plasma cortisol plays a central role in glucose metabolism during the physiological stress response, and it can be used as an indicator of the acute stress response (Wendelaar Bonga, 1997).

The AG group displayed lower cortisol levels than the NAG and DG groups, which, to some extent, corroborated with the efficacy of MS-222 in mitigating the stress of aquaculture. This result was consistent with those of other studies, such as the results obtained after 10 min of enforced exercise in *Scophthalmus maximus* (Van Ham et al., 2003), 5 min of confinement in *Rutilus rutilus* (Pottinger et al., 1999), and handling or confinement stress in *Salmo trutta* (Sumpter et al., 1985) at different temperatures. In contrast, Davis et al. (1984) observed increased plasma corticosteroid levels when the fish were stressed by confinement. These disparities are dependent upon and possibly exacerbated by the species, size, and life experience of the fish as well as the dosage of the anesthetic and type, magnitude, and duration of the stressor (Weber et al., 2011).

In this study, the fish subjected to acute physical stress showed significantly elevated plasma cortisol levels (levels in the NAG group were 1.7-fold those in the CG group). The maximum levels of plasma cortisol in *P. argenteus* were somewhat lower than those in other fishes, such as the Arctic charr (Backström et al., 2017) and *Oncorhynchus kisutch* (Talbot et al., 2009). Therefore, low but significant levels of cortisol after stress could suggest that *P. argenteus* is sensitive to aquaculture treatment stresses and stresses with significant effects on this species. Our results indicate that MS-222 effectively suppresses the cortisol stress response and may prove to be a useful anesthetic for reducing the adverse effects of stress.

### Antioxidant capability

Stressed fish have been demonstrated to be more vulnerable to disease because of impairment of the antioxidant defense systems (Velisek et al., 2011). To cope with oxidative damage, organisms have evolved a system to either prevent or repair the effects of oxidative stress (Birnie-Gauvin et al., 2017). Oxidative stress is caused by the formation of reactive oxygen species (ROS), H2O2, hydroxyl radicals, and superoxide anion radicals, which are the main by-products of oxidative metabolism and potentially cause cell damage (Azzi et al., 2004).

In the present study, the SOD and CAT activities significantly increased (P < 0.05) in the NAG group. This indicates that aquaculture treatment stresses may cause oxidative stress in *P. argenteus* juveniles because SOD and CAT represent the first line of defense against oxidative stress (Farombi et al., 2007). To maintain a balance between antioxidants and ROS, SOD initially converts O^2-^ into O_2_ and H_2_O_2_, and H_2_O_2_ is then broken down into O_2_ and H_2_O by CAT (Li et al., 2009b). CAT is mainly located in the peroxisomes, and it is responsible for the reduction of H2O2 produced by the metabolism of long-chain fatty acids in peroxisomes (Velisek et al., 2011). The SOD and CAT activities significantly decreased (P < 0.05) in the AG group when compared with the NAG group, but significantly increased (P < 0.05) when compared with the CG group. This indicates that MS-222 has a calming effect, but its role is limited. It should be mentioned here that MS-222 significantly reduced MDA content in the liver, indicating that MS-222 could inhibit lipid peroxidation. This was supported by the fact that MDA is a commonly used indicator to evaluate the toxic processes caused by free radicals (Parvez and Raisuddin, 2005). In addition, the NAG group showed significantly increased (P < 0.05) GSH levels when compared with the AG group, but no significant changes were observed when compared with the DG group. Glutathione reductase plays an important role in cellular antioxidant protection and adjustment processes of the metabolic pathways (Wendelaar Bonga, 1997). Glutathione reductase catalyzes the reduction of glutathione disulfide to reduced GSH in an NADPH-dependent reaction (Cazenave et al., 2006). The CAT and GSH activities and MDA contents of the DG group were similar to those of the CG group; however, the SOD activities were significantly increased (P < 0.05) in the DG group. This is probably because drainage is slow and has little influence on oxidative stress. Consistent with those results, In the present study, fish in the NAG group showed more oxidative damage than the control group, which is consistent with this finding; therefore, MS-222 has some functions against this type of damage. This suggests that MS-222 (75 mg L^-1^) can contribute to the health of *P. argenteus* during aquaculture treatments.

### Stress-related gene expression

HSPs, which represent a subset of molecular chaperones, are part of the cellular defense. Within the HSP family, HSP70 and HSP90 play a key role in maintaining cellular homeostasis and protecting the organism after stress(Fu et al., 2011; Liu et al., 2013). Previous studies have shown that HSPs can be regulated by environmental stresses such as heat shock, bacterial challenge, and heavy metals (Farcy et al., 2007) (Ivanina et al., 2009). We observed that the mRNA levels of HSP70 and HSP90 were significantly higher (P < 0.05) in the NAG group than in the other three groups. In addition, no obvious differences in HSP70 and HSP90 mRNA expression levels were observed between the AG and CD groups. When the fish were exposed to stress, over-expression of HSP70 and HSP90 may have been immediately induced to help the cells relieve stress and adapt to the external environmental changes (Ceyhun et al., 2010). This result was consistent with those of the ammonia toxicity test on *Takifugu obscurus* (Cheng et al., 2015) and *Botia reevesae* (Qin et al., 2013). To understand the functions of HSPs, the gene sequences of many marine organisms have been cloned and detected, including the rainbow trout (Ojima et al., 2005), Chinese mitten crab (Li et al., 2009a), black tiger shrimp (Rungrassamee et al., 2010), and silver sea bream (Deane and Woo, 2005). Previous studies have suggested that HSP70 is induced under a certain range of stress, which is a protective mechanism to counteract the effect of stress on cultured animals (Iwama et al., 1998). However, HSP70 in the rainbow trout and tilapia was significantly inhibited under different high-temperature stresses (Basu et al., 2001). A previous study on the Atlantic salmon showed that low temperature, capture, and other stress factors did not significantly influence the HSP levels in the body (Zarate and Bradley, 2003). The reasons for this inconsistency may be related to the type of stress protein. Overall, the mechanisms by which HSPs protect cells under stressful conditions are complex and need to be studied further.

In the AG group, HSP70 and HSP90 mRNA levels were unaffected by stress, whereas GR1 and GR2 mRNA levels were significantly higher (P < 0.05) after stress. This may be because GRs are more sensitive to stress, even after MS-222 administration. In teleosts, two types of GRs (GR1 and GR2) are present, and their functions are affected by the environment and species. For instance, in the Indian ricefish, *Oryzias dancena,* GR1 is believed to be activated only under stressful conditions, and GR2 is activated under non- or mildly stressful conditions (Kim et al., 2011). Sun et al. (Sun et al., 2017) suggested that GR1 is expressed only under stress in *P. argenteus.* In this study, the expression levels of GR1 and GR2 were significantly different between the control and aquaculture treatment groups. Further studies are required to understand the functional model of GRs in this species. In this study, the upregulated mRNA expression levels of HSP70, HSP90, GR1, and GR2 after aquaculture treatment stresses indicate that the HSPs are inducible and may play an important role in the immune response. The mRNA expression levels of HSP70, HSP90, GR1, and GR2 did not differ between the CG and DG groups, indicating that drainage during aquaculture has negligible effects on the expression of stress-related genes.

In summary, MS-222 can be used as an anesthetic for *P. argenteus* juveniles. The most suitable concentration of MS-222 for attaining deep anesthesia was 75 mg L^-1^. An appropriate concentration of MS-222 for sedation can reduce fish activity, facilitate handling procedures, and allow rapid recovery, thereby enhancing the safety of the aquaculture treatment. Aquaculture treatment stresses can cause great damage to *P. argenteus,* and MS-222 (75 mg L^-1^) can help this species to resist the stresses.

## Acknowledgments

This study was funded by The Key projects of the Ningbo people’s livelihood in Agriculture (2013C 11010) and partially funded by Natural Science Foundation of China (31772869), Natural Science Fou ndation of Zhejiang (LY18C190008), Agriculture Key Special Project of Ningbo (2015C110003), Ning bo Livelihood Key Project(2013C11010).The study was also supported by K.C. Wong Magna Fund in Ningbo University, Li Dak Sum Yip Yio Chin Kenneth Li Marine Biopharmaceutical Development Fu nd,National 111 Project of China, and Collaborative Innovation Center for Zhejiang Marine High-Effic iency.

